# Crossmodal correspondence of natural amplitude modulations

**DOI:** 10.1101/2023.09.13.557575

**Authors:** Luis Lemus, Isaac Morán, Jonathan Melchor, Tonatiuh Figueroa, Miguel Matta, Elizabeth Cabrera

## Abstract

Crossmodal correspondence is the ability to recognize objects from different sensory modalities to be similar. Here, we investigated the capacity of humans to perform crossmodal discriminations between visual, acoustic, and tactile stimuli modulated in amplitudes of low-pass filtered envelopes obtained from natural sounds. We found that the perception of crossmodal correspondence emerged from the number of amplitude prominences in each stimulus rather than from temporal variations. Therefore, we propose saliency as the supramodal trade coin of temporal modulations.

## INTRODUCTION

Sometimes, we perceive inputs from different sensory modalities as part of a single object (Gaydos, 1956; Hadjikhani & Roland, 1998; Brown & Brumaghim, 1968). For example, we recognize a buzzing around us and a swiftly flying insect as part of the same entity. However, how our perception achieves such a binding phenomenon still needs to be clarified.

For instance, crossmodal correspondence arises from synchronic stimuli varying in the same supramodal domain, e.g., amplitude, numerosity, duration, and spatial location (Hirsh & Sherrick, 1961; Stein et al., 1988; 1989; Evans & Treisman, 2010; Guzman-Martinez et al., 2012; Takeshima & Gyoba, 2013; Laing et al., 2015; Nieder, 2012); Also, pitch, brightness (Marks, 1989), rhythm, and visual clutter (Sherman et al., 2013). Similarly, accuracy in vocal detection is faster for synchronic visual and auditory cues (Chandrasekaran et al., 2011,2013; Perrodin et al., 2014; Maddox et al., 2015), whereas synchronized but incongruent stimuli lead to illusory percepts like the McGurk effect (McGurk & MacDonald, 1976; Shipley, 1964). Because of the former, it is essential to study crossmodal correspondences among stimuli set apart in time (Stein et al., 2014). Moreover, only a few studies have investigated trimodal correspondences, like the study of Frings and Spence (2010), who found that rhythmicity stimuli facilitated target location.

Experiments on behaving monkeys have found that multisensory brain areas, like the prefrontal, premotor, and ventral intraparietal cortices, represent supramodal information such as numerosity (Nieder, 2012; Vergara et al., 2016). However, numerous studies have recently reported that primary sensory cortices respond to multiple sensory modalities (Zhou & Fuster, 2000). However, it has been argued that multisensory responses in primary cortices might relate to non-parametric processes, such as attention, rather than to conveying information about the physical parameters of stimuli (Lemus et al., 2010).

In the present study, we aimed to study the ability of humans to perform crossmodal discriminations of visual, tactile, and auditory stimuli set apart in time, whose dynamic variations in amplitude resemble those of natural stimuli. Our findings indicate that crossmodal discrimination relies on the number of salient events of each stimulus.

## METHODS

### Subjects and Ethics Statement

Seventeen graduate and undergraduate students from the National Autonomous University of Mexico (UNAM; naïve to the experiment; 8 females, mean age = 25) gave written consent to participate in the psychophysical experiments. All subjects were right-handed and declared no physical impairments. Participants performed 480 trials during a ∼40-minute period but were allowed five-minute breaks at any time. The Bioethical Committee of the Instituto de Fisiología Celular, UNAM, approved the experiments.

### Stimuli

We created acoustic 1 s length (A), visual (V), and tactile (T) stimuli of natural amplitude modulations (NAM) obtained from acoustic envelopes of twenty natural sounds and words (S20) obtained from internet libraries.

#### Acoustic and tactile stimuli

We resampled the S20 to 11025 Hz to homologize. After deriving the envelopes through a Hilbert transform, they were low pass filtered to 15 Hz for smoothing (LP20). The resulting signals were chimerized (Ch20) with white noise as a carrier, normalized to -10 dB RMS, and saved as WAV files (Figure 1A). The sounds were filtered and normalized using Adobe Audition ®. The chimeras were created using MATLAB algorithms previously reported by Smith et al. (2002).

**Figure 1.**
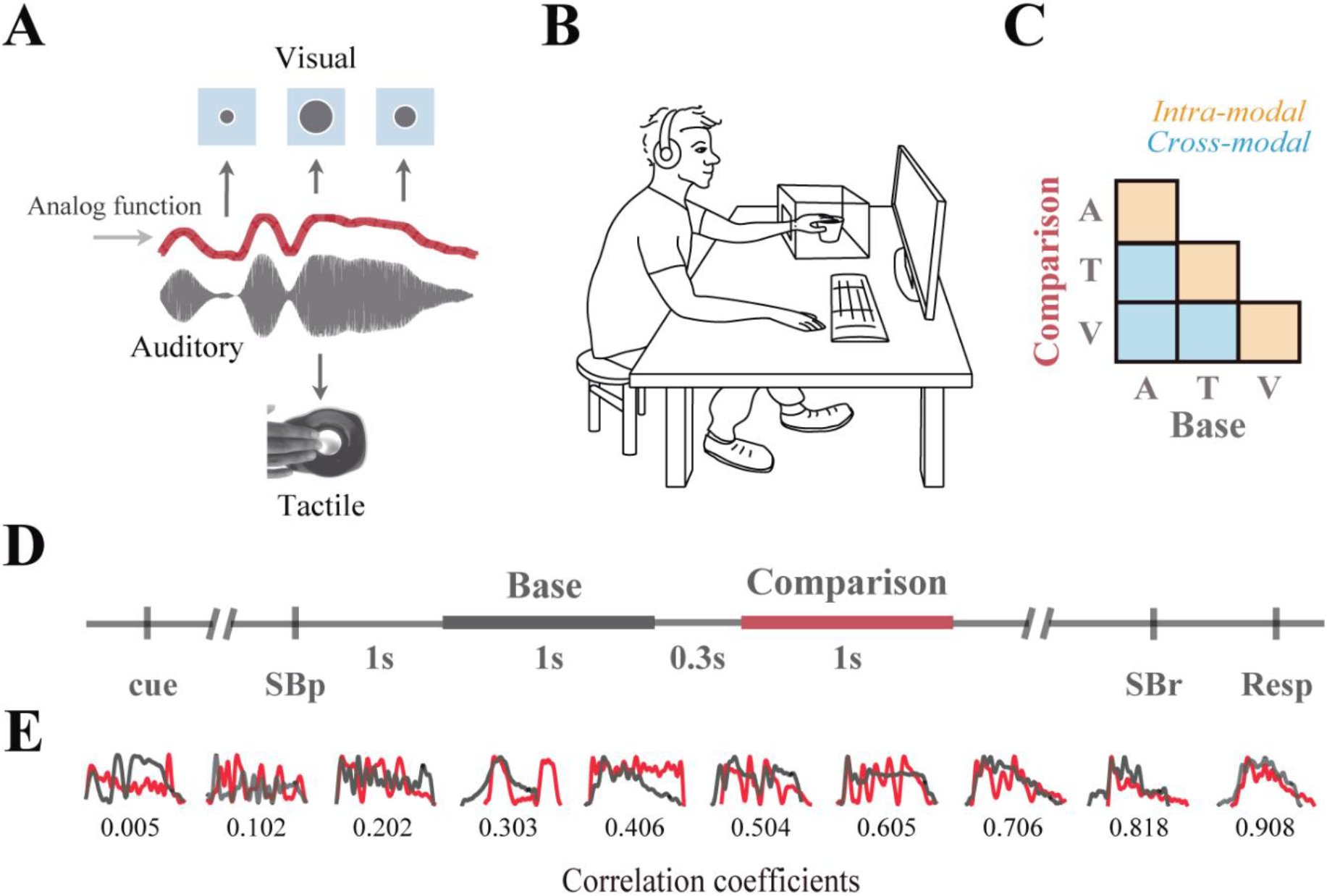
Stimuli and Task. A) Auditory, visual, and tactile stimuli followed the amplitude modulations of diverse acoustic envelopes. The acoustic and tactile stimuli resulted from chimerizing the envelopes with a white noise carrier. The visual stimuli consisted of 1 s AVI clips of a circle of radii varying as the acoustic envelopes. B) Experimental setup. C) Crossmodal and intramodal trials: A, auditory; V, visual; T, tactile. D) Sequence of events in a trial. SBp, spacebar press; SBr, spacebar release; R, response E) Depiction of pairs of stimuli modulations. The first and second stimuli are in black and red, respectively.

#### Visual stimuli

We resampled the (LP20) to the monitor’s 60 Hz frame rate and normalized [1:2]. The sixty resulting values served as radii of JPG circles saved as 1s AVI clips (V20). All stimuli were obtained from lab-customed MATLAB algorithms (The MathWorks Inc).

### Experimental setup

Experiments took place in a psychophysics booth. Each participant sat behind a table facing a 21-inch LCD color monitor (1920 x 1080 resolution, 60Hz refreshing rate) (Figure 1B). The V spanned from 2° to 5° at a 60 cm visual distance. The T stimuli were delivered throughout a speaker inside a homemade soundproof box (Logitech speaker ® Z120 1.2 W, 11.8 cm width x 12.2 cm height; StarTech ® USB sound card) at ∼50 dB SPL. The participants placed their left hand inside the box and touched the speaker’s membrane using the tip of their middle finger. The A monophonic stimuli and white-noise background sounds were delivered binaurally through active noise-blocking headphones (Bose ® QC25) at ∼65 dB SPL and ∼55 dB SPL, respectively. The participants used their right hand for responses on a laptop (Windows 8.1, Intel ® Core i5-3210, 2.5GHz, RAM 6.00GB, 64-bit Operating System).

### Behavioral task

Seventeen human subjects (Power analysis = 0.77) discriminated pairs of intramodal (IM), i.e., VV, TT, and AA, or crossmodal (CM), i.e., AT, AV, and TV (Fig 1C) stimuli presented sequentially in each trial (Figure 1D). Participants commenced a trial by holding the spacebar down after a 2° white cross appeared at the center of the monitor. A second later, a sequence of a first 1s stimulus, a 0.3 s delay, and a second 1s was delivered. Then, subjects released the spacebar and button number 1 if the stimuli were equivalent or button number 2, nonequivalent. The task was programmed in LabVIEW 2014 (SP1 64-bits, National Instruments ®). The statistical analyses were performed using Sigma Plot® version 12.0 for Windows (Systat Inc.).

### Set of trials

We created sets of IM and CM pairs from combinations of Ch20 and V20. Since the task was devised to find whether IM and CM stimuli were perceived as equivalent (E) or nonequivalent (NE), we first estimated the similarity between stimuli pairs by computing the Pearson’s r correlation of the Low-Pass envelopes aligned at the best cross-correlation lag. Afterward, to create each Set, we selected ten pairs of NE from pairs whose r values ranged from ∼0 to ∼0.9 in steps of ∼0.1 and ten E pairs, created by pairing two single stimuli, i.e., r = 1 (Figure 1E). A block of trials consisted of [(3 CM + 3 IM) x Set] = 120.

## RESULTS

We evaluated the ability of human subjects to discriminate NAM stimuli presented in IM and CM trials. In each trial, a subject reported whether a first stimulus was equivalent or nonequivalent to the second stimulus. Crucially, a block of trials comprised pairs of stimuli set in a similarity continuum ranging from 0 to 1 Pearson’s r correlations.

### Uncorrelated crossmodal pairs are often perceived as equivalent

Figure 2A shows the overall CM accuracies (Table 1). For AT condition was 80.7 % ± 4.6 (mean ± s.e.m.), AV 76.9 % ± 6.5, and TV 71.1 % ± 6.7. All distributions were significantly higher than chance at 50% accuracy, T-test, p< 0.001. Specifically, AT: t (16) = 26.9, p < 0.001; AV: t (16) = 16.8, p < 0.001; TV: t (16) = 12.8, p < 0.001. Differences were statistical between AT and TV conditions (rm-ANOVA2 F (11, 176) = 33.5, p < 0.001; post hoc comparisons using the Tukey test for were: AT, p < 0.001; AV, p = 1.00; TV, p < 0.001; Table 1). However, only AT E differed from AT NE (Figure 2B).

**Table 1.**
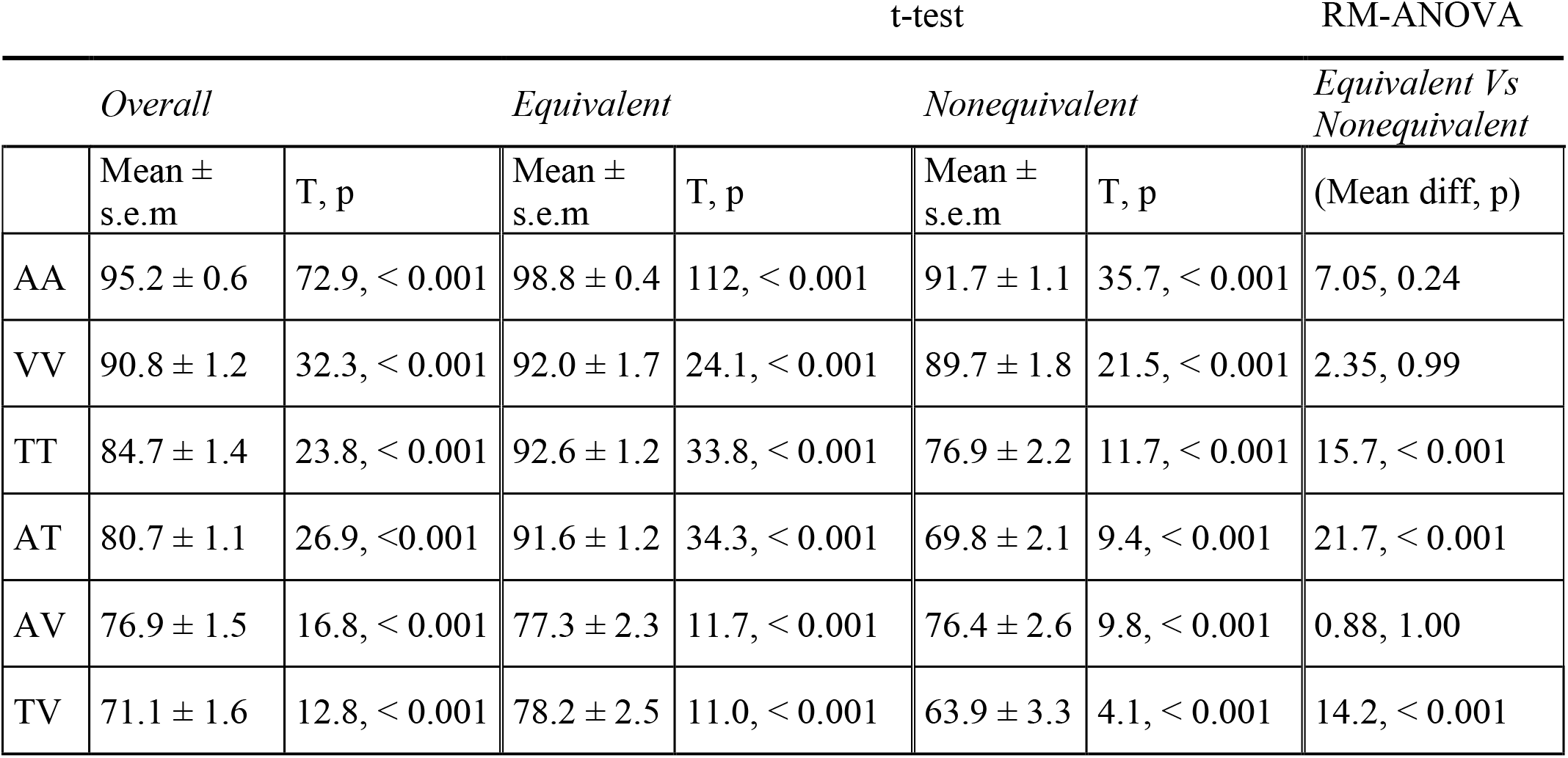
Crossmodal and intramodal accuracies.

**Figure 2.**
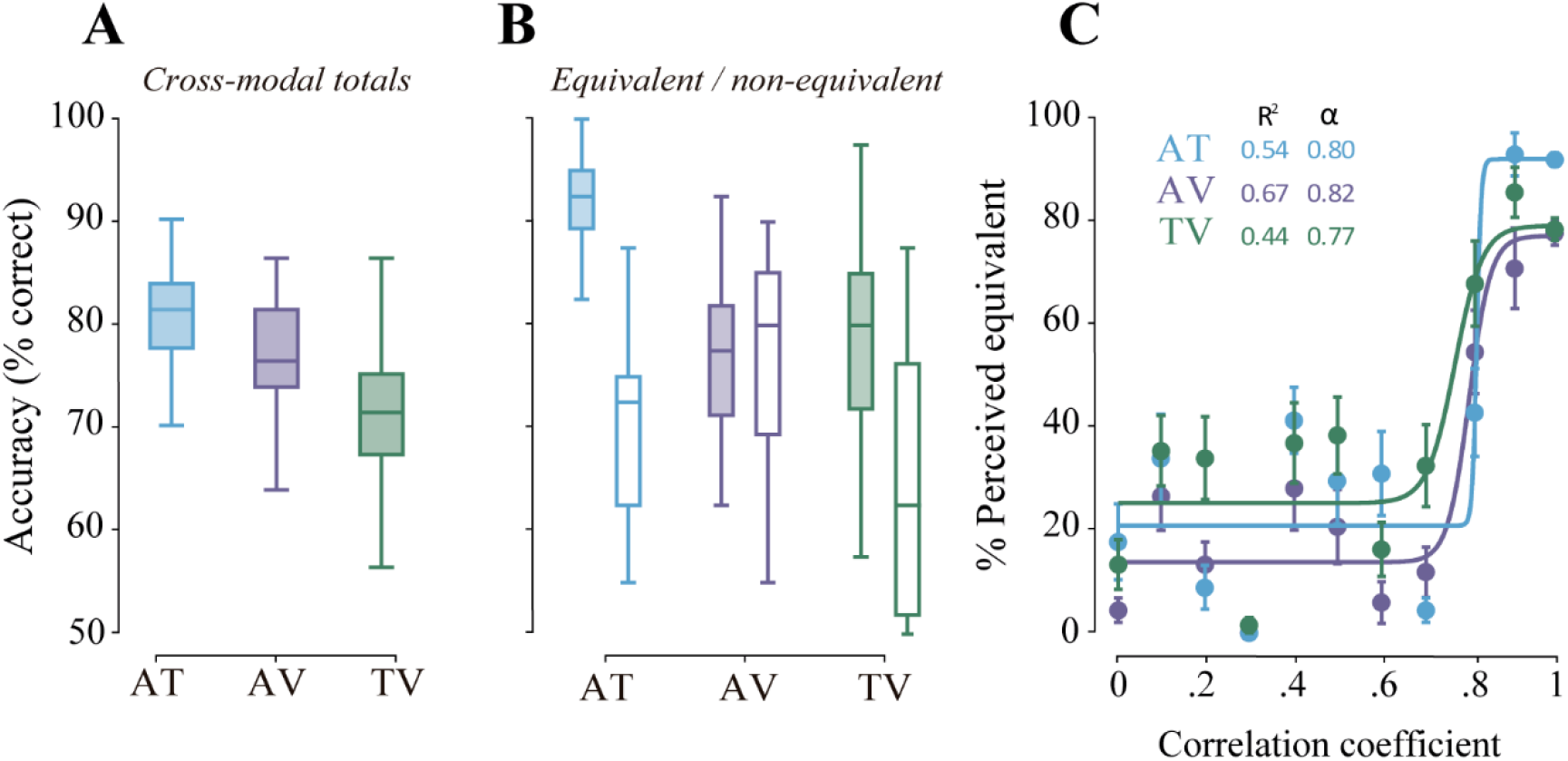
Performance during crossmodal discrimination. A) Boxplots of the participants’ overall performance across crossmodal conditions. AT, audio-tactile; AV, audio-visual; TV, tactile-visual. B) Performance at equivalent and nonequivalent pairs. C) Sigmoidal fits of accuracy as a function of Pearson’s r between the different base and comparison pairs.

Interestingly, the mean AT E distribution was above 90% accuracy, while AT NE was below 70%. This observation shows how subjects were more capable of finding CM equalities than differences. Figure 2C shows that NE pairs below 0.8 r were more inaccurate than the E for all CM conditions. Moreover, the NE-E perceptual thresholds obtained from logistic regressions to the data were 0.80, 0.82, and 0.77 for AT, AV, and TV, respectively. In other words, finding CM differences below those r values was more challenging than finding similarities.

### Intramodal comparisons confirmed a perceptual bias in finding similarities between two stimuli

To determine the extent to which CM discrimination depended on sensory appropriateness. We presented the participants with IM trials, i.e., AA, VV, and TT. Figure 3A shows the overall performance for each IM condition: AA (95.2 ± 2.5 %), VV (90.8 ± 5.2 %), and TT (84.7 ± 6.0 %) (Table 1). A T-test analysis proved that all IM conditions were above chance: AA, t (16) = 72.9, p = < 0.001; VV, t (16) = 32.3, p = < 0.001; TT, t (16) = 23.8, p = < 0.001. Moreover, all E accuracies exceeded NE (Figure 3B), being statistical for TT(rm-ANOVA2 F (11, 176) = 33.5, p < 0.001 post hoc comparisons using the Tukey test (p < 0.001; Table 1).

**Figure 3.**
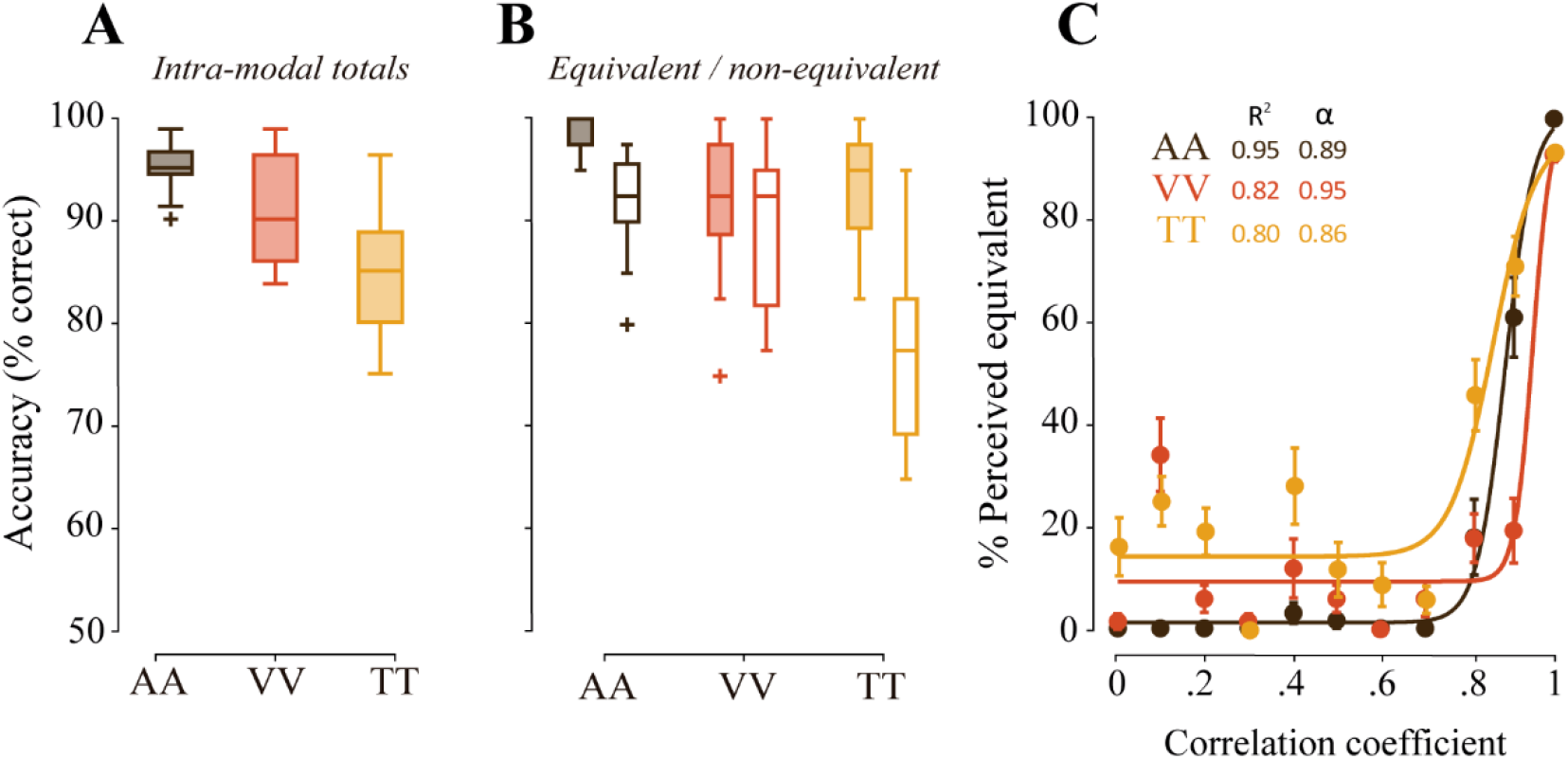
Performance during intramodal discrimination. A) Boxplots of the participants’ overall performance across intramodal conditions. AA, acoustic-acoustic; VV, visual-visual; TT, tactile-tactile. B and C, same as in Figure 2, but for intramodal conditions.

Figure 3C presents the sigmoidal fit to the data. Here, the computed discrimination thresholds were the following: AA (alpha = 0.89, R2 = 0.95), VV (alpha = 0.95, R2 = 0.82), and TT (alpha = 0.86, R2 = 0.80). All IM discriminations resulted better than the CM. Noticeably, even the worst IM performance, i.e., TT, was better than the best CM performance, i.e., AT. We interpret this result as CM suffers from a loss of information during the discrimination process rather than from appropriateness differences in sensory modalities. Furthermore, given that both IM and CM discriminations showed differences between E and NE, we conclude that the discrimination of two NAM stimuli favors finding similitudes rather than discrepancies.

## DISCUSSION

In the present study, we asked human subjects to report whether two consecutive NAM stimuli delivered intra, or crossmodally were perceived as equivalent or nonequivalent. We set stimuli apart for preventing multisensory fusion effects such as those observed during superimposed stimuli (McGurk & McDonald, 1976; Shipley, 1964; Parise & Ernst, 2016; Bizley et al., 2016), but only 0.6 s to reduce informational loss in working memory. Most IM and CM discrimination thresholds were above ∼0.8 r, which was consistent with other studies showing that CM correspondence increased as a function of the mutual coherence of stimuli (Frings & Spence, 2010; Parise et al., 2012; 2013; Guzman-Martinez et al., 2012; Takeshima & Gyoba, 2013; Chandrasekaran et al., 2011;2013; Maddox et al., 2015). However, we found that many times NE stimuli (i.e., < 0.8 r) were perceived as equivalent (Wozny et al., 2008; Bresciani et al., 2008).

Since IM false alarms were less frequent than CM, and given that our task did not involve working memory, IM discriminations probably took place at primary and secondary sensory cortices (Fuster et al., 2000; Romo et al., 2002; Lemus et al., 2009a; 2010) and not necessarily at polysensory areas, like in paradigms requiring working memory (Man et al., 2015; Ungerleider et al., 1998; Romo et al., 1999; Lemus et., al 2009b; Calvert, 2001; Man et al., 2015; Nieder, 2012; Vergara et al., 2016).

Another finding in our experiments was that the participants accurately perceived 0.3 r as NE for all CM and IM conditions. Figure 1E shows that 0.3 r consisted of stimuli of one and two salient events, respectively. Therefore, subjects discriminated at those trials better than those of higher r values. An inference is that the subjects discriminated the number of saliencies rather than overall correlations between stimuli, which is consistent with neuronal models of crossmodal comparisons (Bizley et al., 2016; Parise & Ernst, 2016). Future experiments may elucidate the role of saliency in the recognition of objects.

## CONCLUSIONS

The results show that the perception of crossmodal correspondence between auditory, tactile, and visual stimuli occurs in pairs of poor correlations, where only a few prominent signals can be categorized. Therefore, we predict that crossmodal comparisons at polysensory cortices rely on saliency rather than the fine structure of stimuli.

## Conflict of Interest

The authors declare no conflict of interest.

## Funding

This work was supported by the Consejo Nacional de Ciencia y Tecnología (CONACYT) 181301. El Programa de Apoyos a Proyectos de Investigación e Innovación Tecnológica (PAPIIT) IA201513.

## Acknowledgments

We thank Centenario 107 and its crew for the valuable time, space, and edible resources. We also value the support of the Computer Unit, Ana Escalante, and Francisco Pérez of the Instituto de Fisiología Celular, UNAM.

## Notes

### Competing Interest Statement

The authors have declared no competing interest.

